# System analysis of differentially expressed miRNAs in hexaploid wheat display tissue-specific regulatory role during Fe deficiency response

**DOI:** 10.1101/2023.02.19.529099

**Authors:** Shivani Sharma, Dalwinder Singh, Riya Joon, Vishnu Shukla, Ajit Pal Singh, Palvinder Singh, Shrikant Mantri, Ajay K Pandey

## Abstract

**Background:** Iron (Fe) is an essential mineral element, and its deficiency in soil largely affects crop productivity. In plants, the molecular mechanisms underlying the genetic regulation of Fe deficiency responses have yet to be well understood. Specifically, microRNA (miRNA) mediated regulation of Fe deficiency response and its regulatory network is largely elusive. In the current work, we utilized a whole genome transcriptomic approach to identify the Fe deficiency-responsive miRNAs to understand the molecular mechanisms of Fe deficiency response in wheat seedlings. The study also identifies nine novel miRNAs putatively involved in Fe deficiency response. Further, the identified miRNAs showed tissue preferences relating them to differential mechanisms against Fe deficiency.

**Results:** In the present study, we performed small RNA-targeted whole genome transcriptome analysis to identify the involvement of sRNAs in Fe deficiency response. The analysis identified 105 differentially expressed miRNAs corresponding to Fe deficiency response, among them, 9 miRNAs were found to be novel in this study. Interestingly, tissue-specific regulation of Fe deficiency response also participates through miRNA-mediated regulation. We identified 17 shoot specific miRNAs and 18 root-specific miRNAs with altered expression. We validated the tissue specificity of these *miRNAs* by stem-loop quantitative RT-PCR. Further, an attempt was made to predict their targets to speculate their participation in Fe deficiency response. This miRNA target prediction analysis suggested a few major targets of the identified miRNAs, such as multicopper oxidases, E3 ubiquitin ligases, GRAS family, and WRKY transcription factors previously known to play key roles in Fe homeostasis. Our analysis of selected miRNAs also confirmed a temporal regulation of the response.

**Conclusion:** The first information generated here will classify the repository of wheat *miRNAs* (with few novel miRNAs) for their role in Fe deficiency response. Our work provides insights into miRNA-mediated regulatory pathways during Fe deficiency.

## Background

Iron (Fe) is an essential micronutrient. Being the principal component of chlorophyll, Fe-S clusters of enzymes and cofactors it participates in various biochemical processes, including photosynthesis, respiration etc. [1]. Despite being the fourth most abundant element in the earth’s crust, its bioavailability to plants is restricted owing to its presence as sparingly soluble Fe^3+^ form in aerobic and high pH soil environment [2,3]. Fe deficiency in plants causes interveinal chlorosis and drastically impacts vegetative growth and crop yield [4]. Therefore, plants have devised distinct uptake mechanisms and efficient modes of Fe translocation for its tissue-specific distribution via specific transporters and chelators [5]. Dicots and non-graminaceous monocots utilise the strategy-I mode of Fe uptake by reduction of Fe^3+^ to Fe^2+^ at the root surface, followed by internalization of soluble Fe by membrane-localized iron-regulated transporter-1 (IRT1) [6]. Graminaceous plants like maize and wheat, on the other hand, chelate Fe^3+^ by excreting phytosiderophores, chelation-based strategy known as strategy-II. The resulting complexes are then taken up by yellow stripe-like (YSL) transporter in the plasma membrane of roots. Molecular components regulating Fe homeostasis in plants comprise distinct families of transcription factors (TFs) like basic helix-loop-helix (bHLH), WRKY and members of no apical meristem (NAC), IDE-binding factors (IDEF1 and IDEF2) [7–13]. Some of these TFs are regulated at post-translational levels by E3-ubiquitin ligases such as BRUTUS (BTS) in Arabidopsis and Hemerythrin-rich zinc finger-like proteins (HRZ1 & HRZ2) in rice to tackle the deleterious consequences of activation of Fe uptake and transport machinery during Fe deficiency [14,15]. The studies related to Fe transport and regulation have been extensively carried out in model plants like Arabidopsis and rice but to only a limited extent in wheat.

Plant responses are regulated by a number of non-coding RNAs (ncRNAs) where these ncRNAs can be categorised into three major classes i.e. small (18-30 nt), medium (31-200 nt) and long (>201 nt) based on their nucleotide length [16]. MicroRNAs (miRNAs) are a single-stranded non-coding endogenous class of small RNAs that regulate the target mRNAs by either causing their cleavage or translational repression [17]. In plants, *miRNAs* are known to play roles in diverse fundamental processes as controllers of vegetative and floral organ development, phytohormone signalling, and regulation of various biotic and abiotic stress responses, including regulation of genes involved in nutrient uptake and transport under nutrient stress conditions [18–26]. Previously, 24 miRNAs were found to have iron-deficiency responsive cis-elements (IDE1 and IDE2) in their promoter regions. Around 70% of them i.e. 17 miRNAs were responsive to Fe deficiency in Arabidopsis shoot and/or root [27]. In another study, 32 Fe deficiency responsive miRNAs were identified using a microarray-based approach in leaves, roots and nodules of common bean (*Phaseolus vulgaris*) [28]. Differential expression pattern of miRNAs responsive to Fe deficiency in Arabidopsis rosette and shoot has also been analysed, pinpointing eight miRNAs from seven different families [29]. Recently, several seed iron concentration-related QTLs were found to be the targets of Fe deficiency responsive miRNAs in rice recombinant inbred lines (RILs) [30]. Seven of nine miRNAs identified in the study showed downregulation in response to Fe-deficient conditions. Further, identifying 26 known and 55 novel Fe-deficiency-responsive *miRNAs* in *Citrus sinensis* suggested a larger role being played by miRNAs during Fe deficiency [31].

Hexaploid wheat (*Triticum aestivum* L.) is the most widely grown cereal crop in many countries and accounts for a total of 20% of calorific intake by humans [32]. Multiple miRNAs from wheat have already been characterized for their roles in various abiotic stresses, including nutrient starvation (nitrogen and phosphate), salinity and drought stress [33–35]. Wheat has a complex genetic architecture, and there is a dearth of knowledge about different means of genetic regulation of Fe homeostasis in this crop. Identifying distinct *miRNAs* and their targets during Fe-homeostasis will help develop regulatory networks. Previously, core components participating in strategy-II mode of Fe uptake and mobilization were identified using transcriptomics-based approaches in wheat [36–38]. However, information on the role of miRNAs in the Fe deficiency response in wheat is lacking. Therefore, we extended this work to get an insight into the miRNA-based regulation during Fe-deficient conditions. In the current study, we investigated the differentially expressed miRNAs in response to Fe deficiency to understand small RNA-mediated regulation of Fe-homeostasis. Our work identified a sub-set of the shoot and root-specific miRNAs targeting Fe-mobilization in a tissue-specific manner.

## Results

### Analysis of wheat sRNA during Fe deficiency

To get an insight into sRNA (small RNA) mediated regulation of Fe deficiency response in wheat, we performed whole genome sRNA sequencing from the root and shoot tissues of seedlings subjected to Fe deficiency for different time points. As the study aimed to generate a Fe deficiency responsive sRNA inventory, sequencing of sRNA was done with the pooled RNA samples of the respective wheat tissues exposed to Fe deficiency for different time points (**Figure S1**). A total of 14.54 million and 14.46 million reads were obtained for iron sufficient (+Fe), and iron deficient (–Fe) shoots, respectively, whereas 14.97 million and 16.03 million reads were obtained for +Fe and –Fe root libraries. After removing low-quality reads, 14.31 million and 13.72 million clean reads yielded for +Fe and –Fe shoot, whereas 14.77 million and 15.77 million for +Fe and –Fe root, respectively. After further refining the sequence reads, we ended with 6.4 million to 10.01 million total sRNA reads with 1.74 million to 2.28 million unique reads encompassing different types of small RNAs (**Table S1**). Chromosomal distribution of the mapped sRNA reads on the wheat reference genome showed a predominant contribution from A and B genomes compared to D genomes (**Figure S2**). Interestingly, all the genomes of chromosome 7 contributed to the total sRNA reads.

Interestingly, we observed that the sense strand of the genome is more responsive towards Fe deficiency as more than 61% of the sRNA reads observed were coded from the sense strand, while the antisense strand was only able to contribute for around 5-10% of the fraction, irrespective of the tissue (**Table S1; Figure S2**). Further, we observed a decrease in sRNA reads coded by both sense and antisense strands in the shoot in response to Fe deficiency (4.46% and 1.03% decrease for sense and antisense strands, respectively). In contrast, to shoot, there was an increase in sRNA reads in roots in response to Fe deficiency, preferably over-representing sRNAs coded by antisense strand (0.2% and 1.48% for sense and an antisense strand, respectively).

To gain an insight into sRNA distribution in the genome, we analyzed whether the coding region or the non-coding region is responsive to sRNA-mediated regulation of Fe deficiency response. Further, the positional mapping of generated sRNA reads from the root and shoot accounted for 11.15 % and 6.24 % of reads mapping to exon and 2.54 % and 3.90 % to intron, respectively (**Table S2**).

### Identification of Fe deficiency-induced miRNAs

To characterize the sRNA reads into different subfamilies, we annotated all the sRNAs with Rfam database into rRNA, snoRNA, snRNA, TAS, and miRNA classes (**Figure 1A-D**). This analysis extended our observation of inverse relations in sRNA reads in shoot and root tissues in response to Fe deficiency. This categorised data showed a significant decrease in miRNA representation in response to Fe deficiency in shoot tissues. In contrast, we observed increased miRNA reads in response to Fe deficiency in root tissues (**Figure 1A-D**). This data suggested that miRNAs might be acting in a tissue manner in regulating Fe deficiency response in wheat.

**Figure 1:**
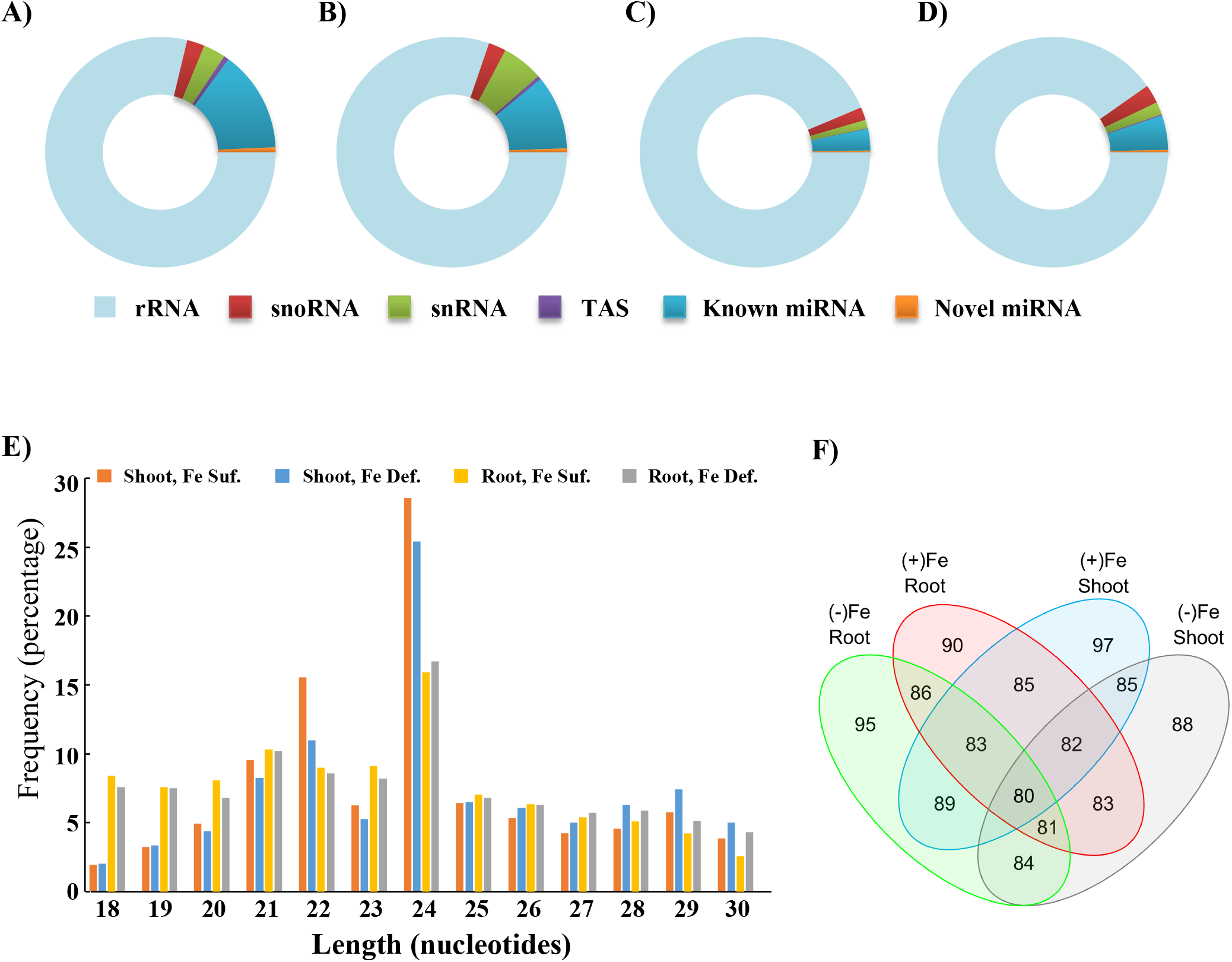
Expression of Fe-regulated sRNAs in hexaploid wheat tissues. Doughnut pie chart showing a differential abundance of sRNAs families in wheat **A)** Fe sufficient shoot, **B)** Fe deficient shoot, **C)** Fe sufficient root and **D)** Fe deficient root tissues. **E)** The total length distribution of Fe-deficiency responsive sRNAs identified from *Triticum aestivum*. The abscissa is the length of sRNAs reads, the ordinate is the percentage of one length read accounted for total sRNAs. **F)** Venn diagram showing comparative analysis of differentially expressed miRNAs in Fe-responsive root and shoot sRNA libraries.

While analysing the length-based classification of unique sRNA reads, we found that the 20-24 nucleotide sRNAs were the most abundant classes in our datasets, representing around 5-30% of our data irrespective of the treatment conditions (Figure 1E). To our interest, the abundance pattern of sRNAs observed in shoot and root tissues w.r.t. to Fe deficiency was also found to be length biased. We observed that 24 nucleotide-long sRNAs follow decreased abundance in shoot while showing an increased abundance in root tissue in response to Fe deficiency (**Figure 1E**).

As we observed that the 20-24 nucleotide long sRNAs are over-represented in our dataset, which typically lies in the range of miRNAs, we analysed these reads for characteristics of miRNAs. Our analysis identified 105 miRNAs in our transcriptome analysis, out of which nine were novel (**Table S3**). As evident from previous reports, most active miRNAs prefer U at the first nucleotide at 5’ end, which in addition to high A+U content, provides them low internal stability, promoting them to be processed into mature miRNA through RISC complex (RNA-induced silencing complex). Additionally, A or U at the 10^th^ position is over-represented in natural plant miRNAs, further contributing to their processibility [39,40]. In agreement of these previous reports, we observed that around 27% of the identified miRNAs showed first nucleotide preference for U and around 45% of miRNAs have nucleotide preference for A/U at 10^th^ position (**Table S3**). Hairpin analysis of these identified miRNAs classified them into 36 miRNA encoding families. Interestingly, members for 35 out of 36 hairpin families are represented in the wheat genome, which was the maximum variability observed among all the 66 plant species analysed (**Table S4**). To further validate our report, we predicted the secondary structure of the identified miRNAs with an RNAfold web server with a minimum free energy index (MFEI) algorithm (http://rna.tbi.univie.ac.at//cgi-bin/RNAWebSuite/RNAfold.cgi). Characteristic stem-loop hairpin formation in all 9 novel miRNAs validated their secondary structure and strengthened their putative function as miRNAs (**Figure S3**).

Next, to pinpoint miRNAs responsive to Fe deficiency, miRNAs were profiled for differential expression patterns using DEGseq in all four libraries. Our analysis revealed that 23 miRNAs were upregulated in the shoot; 17 were shoot specific. Further, 20 miRNAs showed downregulation, with 10 miRNAs downregulated in a shoot-specific manner (**Figure 2**). We extended a similar analysis for root tissues where we observed that 15 miRNAs showed upregulation and 19 miRNAs showed downregulation with 8 and 10 miRNAs with root-specific behaviour (**Figure 2**). This relative analysis further allowed us to identify the miRNAs with similar regulation irrespective of the tissues. We found 3 miRNAs (*tae-miR1122b-3p, -1138* and *-9652-5p*) commonly upregulated while 6 miRNAs (*tae-miRNA5049-3p, novel_1, 395b, 395a, 408* and *9666b-3p*) commonly downregulated in either tissue in response to Fe deficiency. Further, 7 miRNAs (*tae-miRNA9679-5p, -9782, -1118*, - *1117*, -*9674a-5p*, -*9654a-3p* and *novel_10*) showed inverse behaviour in terms of their transcriptional induction during Fe deficiency when shoot and root tissues are compared (**Figure 2**).

**Figure 2.**
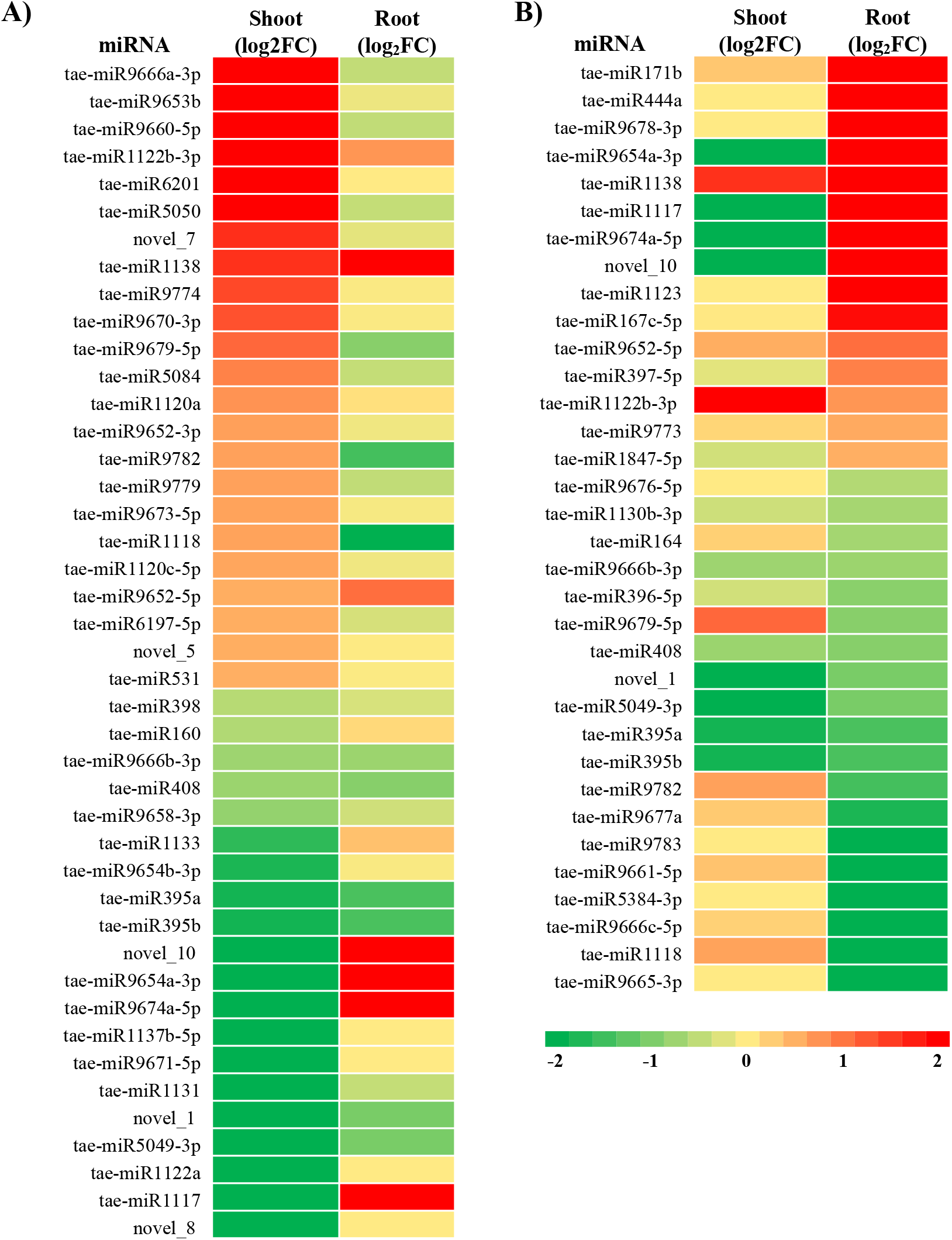
Expression of Fe-regulated miRNAs in hexaploid wheat tissues. Heat map showing differentially expressed miRNAs in shoot and root. A) Significantly up and down-regulated miRNAs in the shoot with their expression values in root in response to Fe deficiency. B) Significantly up and down-regulated miRNAs in root with their expression values in the shoot. Heat maps were plotted against the log2FC values of each miRNA in response to Fe deficiency. FC=1.5 was considered as the criteria for significance.

### Analysis of miRNA for their temporal expression responses

In order to validate our transcriptomic results, we checked the expression of miRNAs in both the tissues. We randomly selected eight miRNAs from our transcriptomic data and performed stem-loop qRT-PCR analysis with gene-specific primer sets (**Table S5**). Following this analysis, we found that the expression pattern of all the selected miRNAs was very similar to the one observed in transcriptomic analysis with Pearson’s correlation coefficient (R^2^=0.9803) (**Figure 3**).

**Figure 3.**
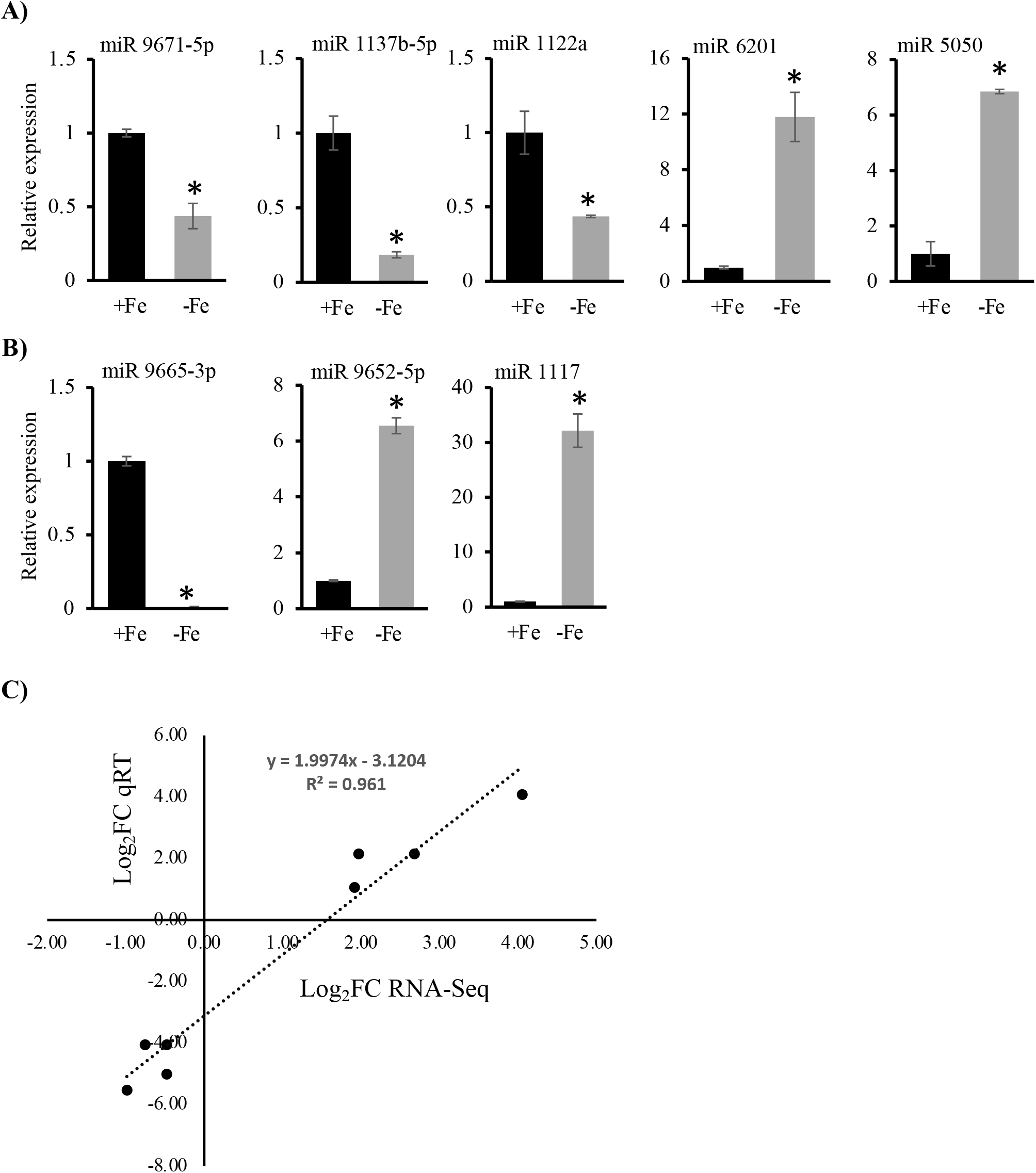
Stem-loop qRT-PCR-based validation of transcriptome data. Expression profile of selected miRNAs in **A)** shoot **B)** root in response to Fe deficiency in wheat. **C)** Line graph showing correlation of qRT-PCR based analysis with transcriptome expression of selected miRNAs in shoot and root tissues of wheat in Fe deficiency response. Relative fold change in the expression was calculated (n=3) after performing the qRT PCR analysis. Two-tailed student’s *t*-test (p<= 0.05) was used to determine the significant change.

To further understand miRNA-mediated Fe deficiency response temporally, we studied expression patterns of multiple miRNAs in either of the tissues after subjecting them to Fe deficiency stress for 6, 9, 12 and 15 days. We selected miRNAs found differentially regulated in the wheat shoot (*tae-miR6201, -5050, -9774, -1122a, -1137b-5p* and *-9671-5p*) and root (*tae-miR1138*, -*167c-5p*, -444a, -*9652-5p*, -*9654a-3p* and -*397-5p*) tissues. Although we found a perfect correlation between our transcriptome and stem-loop qRT-PCR data, varying temporal transcriptional responses of miRNAs were observed during early and prolonged Fe deficiency stress, irrespective of the tissue considered (**Figure 4**). This approach helped us to characterize the tissue-specific expression responses into three categories. The first includes early responsive miRNAs, where most of the miRNAs showed early induction in the shoot, though, in the root, few miRNAs showed a low early response (*tae-miR444a, -9654a-3p* and -*397-5p*). The second category comprises late responsive miRNAs mainly accumulated in root tissues, including (*tae-miR444a, -167c-5p*, -*9654a-3p* and -*397-5p*). Additionally, in shoot tissues, most of them showed weak late responsive nature except two miRNAs (*tae-miR1137b-5p* and *tae-miR9671-5p*) which showed constantly induced and late responsive behaviour, respectively, in response to Fe deficiency (**Figure 4**). The third category highlights the miRNAs with mixed temporal expression during Fe deficiency, which included *tae-miR9774* in shoot and *tae-miR1138* and *tae-miR-9652-5p* in the root. Altogether, our study observed a time-dependent regulation of the wheat miRNAs and potentially established the molecular responses during Fe deficiency.

**Figure 4.**
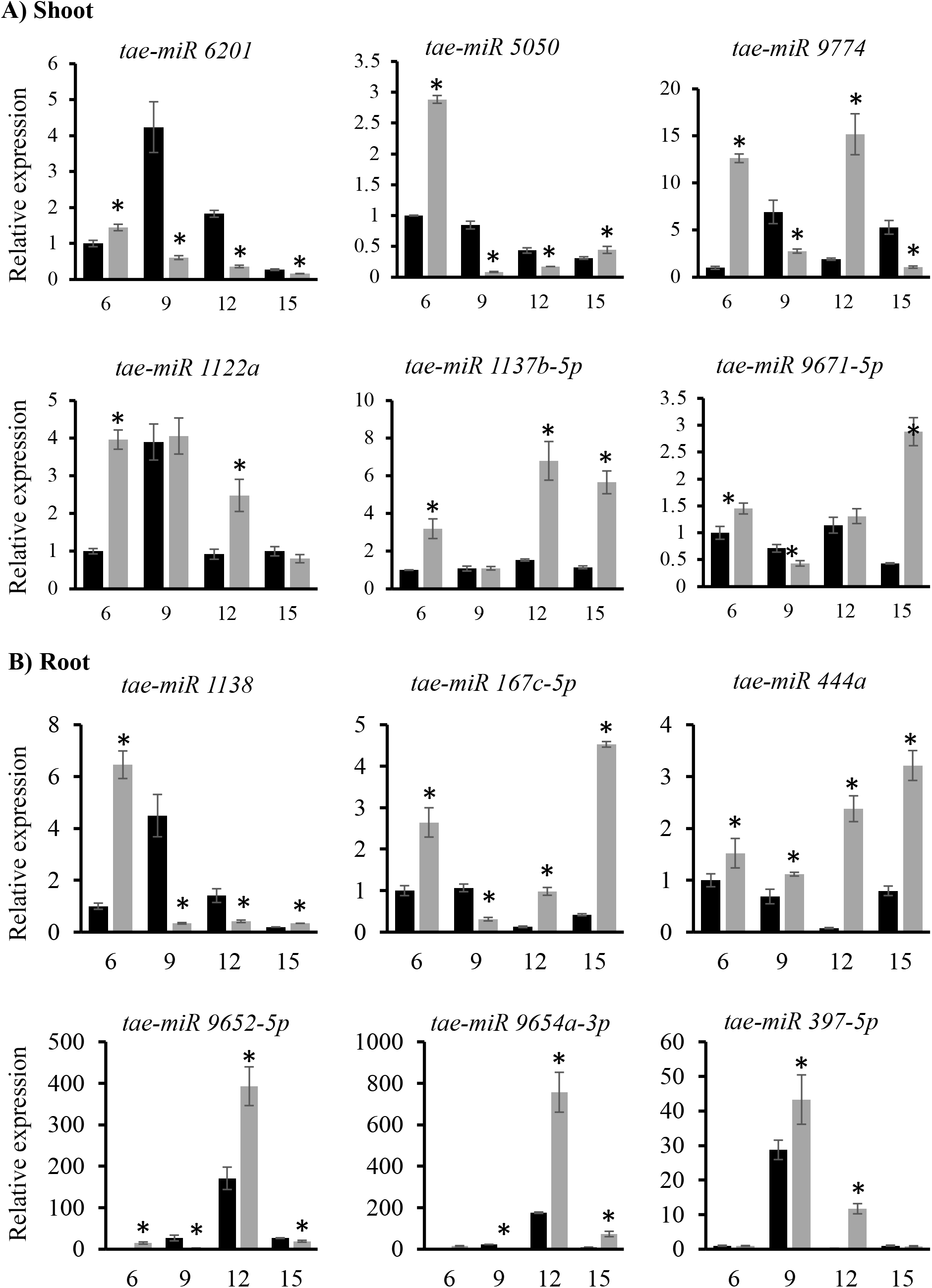
Temporal expression profiling of selected Fe deficiency responsive miRNAs in wheat. Expression profile of selected miRNAs in **A)** shoot **B)** root in response to Fe deficiency in wheat. Relative fold change in the expression was calculated (n=3) after performing the qRT PCR analysis. Two-tailed student’s *t*-test (p<= 0.05) was used to determine the significant change.

### Sub-genomic expression of miRNAs in wheat

In order to further comment on the involvement of different genomes of wheat in the regulation of miRNA expression, we analysed the expression of the differentially expressed miRNAs in in-silico expression analysis with PmiRExAt database (http://pmirexat.nabi.res.in/). The database provided us with the expression values for 62 out of 105 differentially expressed miRNAs in *T. aestivum* (AABBDD), *T. durum* c.v. Langdon TTR16 (AABB) and *Aegilops tauschii*, TQ113 (DD) genomes (**Figure S4A**). Our analysis suggested that the DD genome progenitor *A. tauschii* expressed the least number of miRNAs (27%) while the incorporation of the DD genome into AABB genome (*T. durum*) only partially increased the number of miRNAs expressed (**Figure S4B**). Therefore, though we did not find any negative impact of the DD genome, the AABB genome contributed most to the expression of the selected miRNAs. Although, when we analysed the expression levels of selected miRNAs in different genomes, the AABBDD genome contributed the most to the expression levels of miRNAs (48%), while the AABB genome was the least contributor (21%) while the DD genome contributed around 31% for the expression levels of the miRNAs (**Figure S4C**). Therefore, in conclusion, we observed that the DD genome has the lowest penetrance in miRNA expression levels and the highest expressivity.

### KEGG and GO enrichment analysis

Next, we performed the KEGG enrichment and Gene Ontology (GO) analysis to predict the biological functions of miRNA target genes. This was done to identify the molecular pathways or processes that could be affected by differentially expressed miRNAs under Fe deficiency in a tissue-specific manner. Firstly, the target genes of differentially expressed microRNAs were subjected to KEGG enrichment to identify significantly enriched metabolic pathways. Most of the target genes in both root and shoot were significantly enriched in the biosynthesis of secondary metabolites (**Figure 5**). However, some highly significant enrichment was found to be specific in shoots and roots. For instance, metabolic pathways such as ubiquitin-mediated proteolysis, Carbon metabolism, steroid biosynthesis and RNA degradation and were enriched in shoots (**Figure 5**). In contrast, taurine metabolism, fatty acid metabolism, glutamate metabolism, Nicotinate and nicotinamide metabolism, inositol phosphate metabolism, ABC transporters, RNA transport and homologous recombination, were significantly enriched (**Figure 5**) in roots.

**Figure 5:**
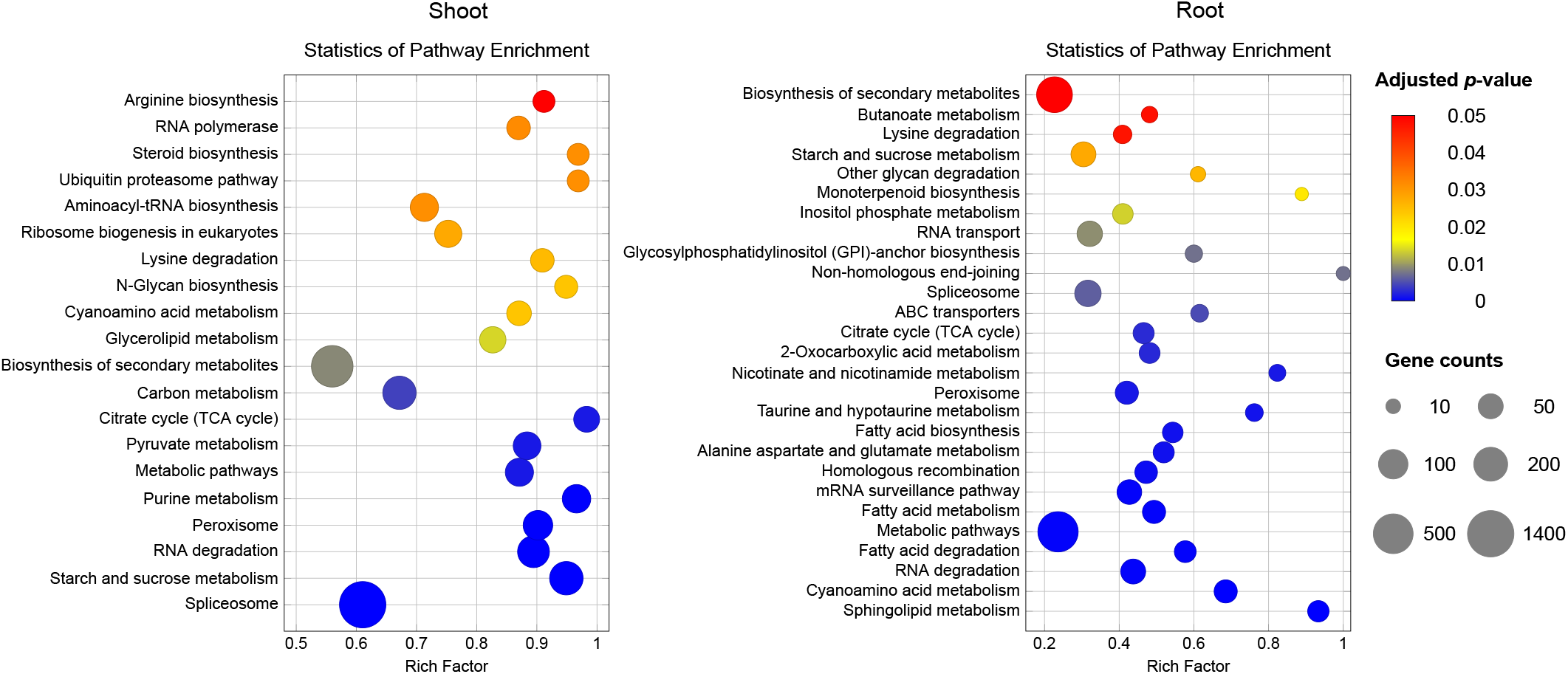
KEGG enrichment for target genes of Fe-responsive miRNAs in the root and shoot. KEGG pathway is displayed along the Y-axis, and X-axis represents all genes enriched in a particular pathway. The colour of the circle indicates the q-value, and the size of the dot correlates with the number of DEGs mapped to the specific pathway.

GO enrichment analysis suggested that both in shoot and root, miRNA target genes are significantly involved in retrograde transport, RNA polymerase complex and DNA damage/conformation change/duplex unwinding/helicase activity (**Figure 6**). Biological processes (BP) such as siRNA processing, triterpenoid metabolism, cell morphogenesis/cell shape regulation, vacuole organization, and protein translation were enriched specifically in shoots. In contrast, target genes in roots were enriched in BP such as prenylated protein catabolism, regulation of abiotic stress, amino acid catabolism and potassium ion membrane transport. Specific enrichment of GO terms belonging to cellular component (CC) was high in shoots. For instance, shoot target genes were significantly involved in GARP complex, myosin complex, mediator complex, ATPase complex and condensed chromosome. In roots, the mitochondrial matrix component was explicitly enriched. Besides, molecular function (MF) such as syntaxin binding, nuclear pore structure constituent, demethylase activity and small GTPase binding were found to be predominant in shoots and enrichment of MF activities mainly, prenylcysteine oxidase activity, glutathione oxidase activity, SNARE binding and molecular adaptor activity were specific to roots (**Figure 7**). Collectively, under Fe deficiency, miRNAs can significantly affect retrograde transport and DNA activities, along with they could target specific biological functions in shoots and roots. We conclude that functional pathway genes in the GO process can overlap in different tissue and be targeted by different miRNAs expressed in a tissue-specific manner.

**Figure 6:**
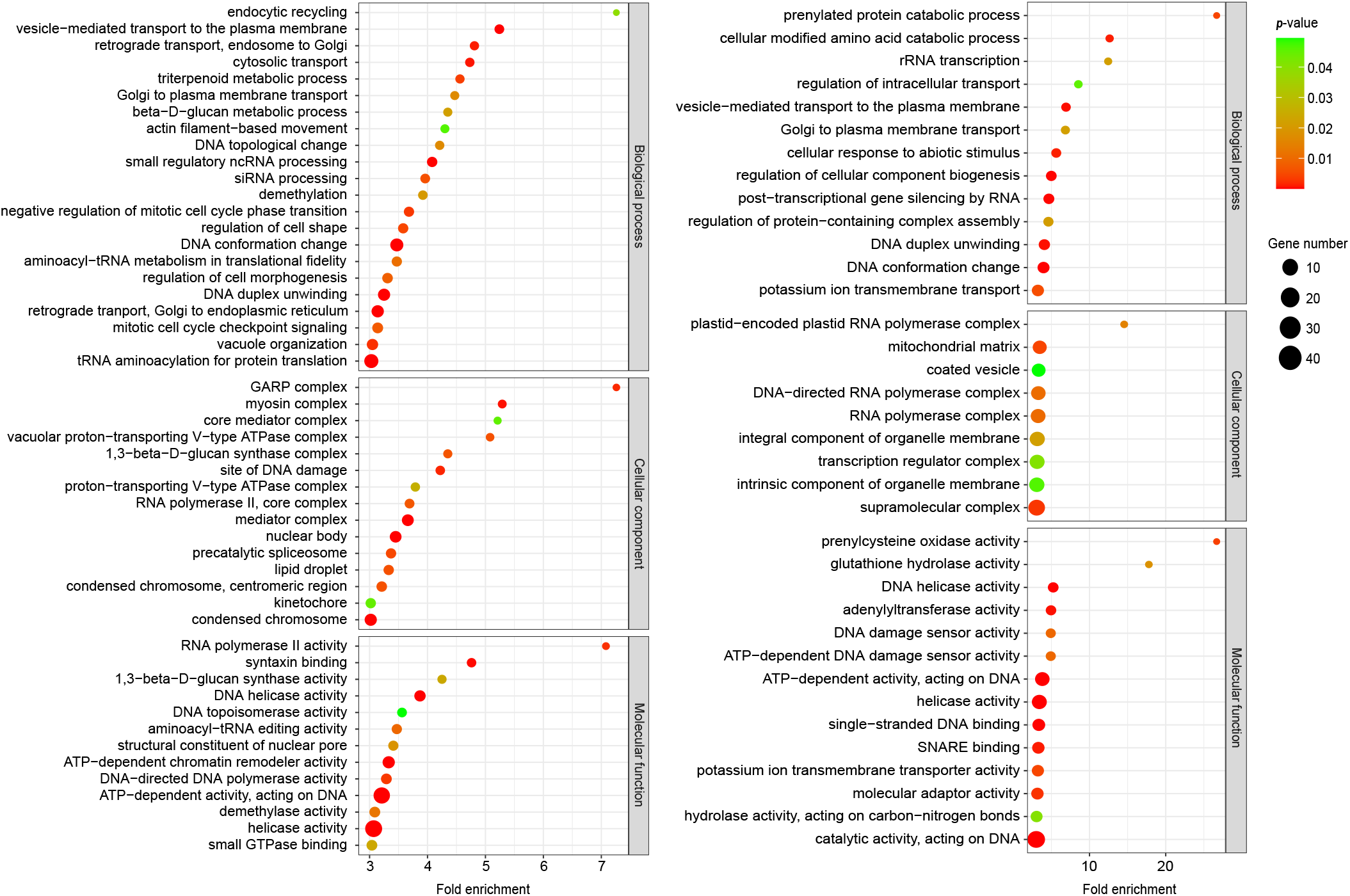
Gene Ontology (GO) categorization of targets of differential expressed Fe-deficiency responsive miRNAs. The y-axis represents the category of miRNA targets, and the x-axis shows. The bubble size indicates the number of genes, with the colour representing the significance as denoted by the *p*-value.

**Figure 7.**
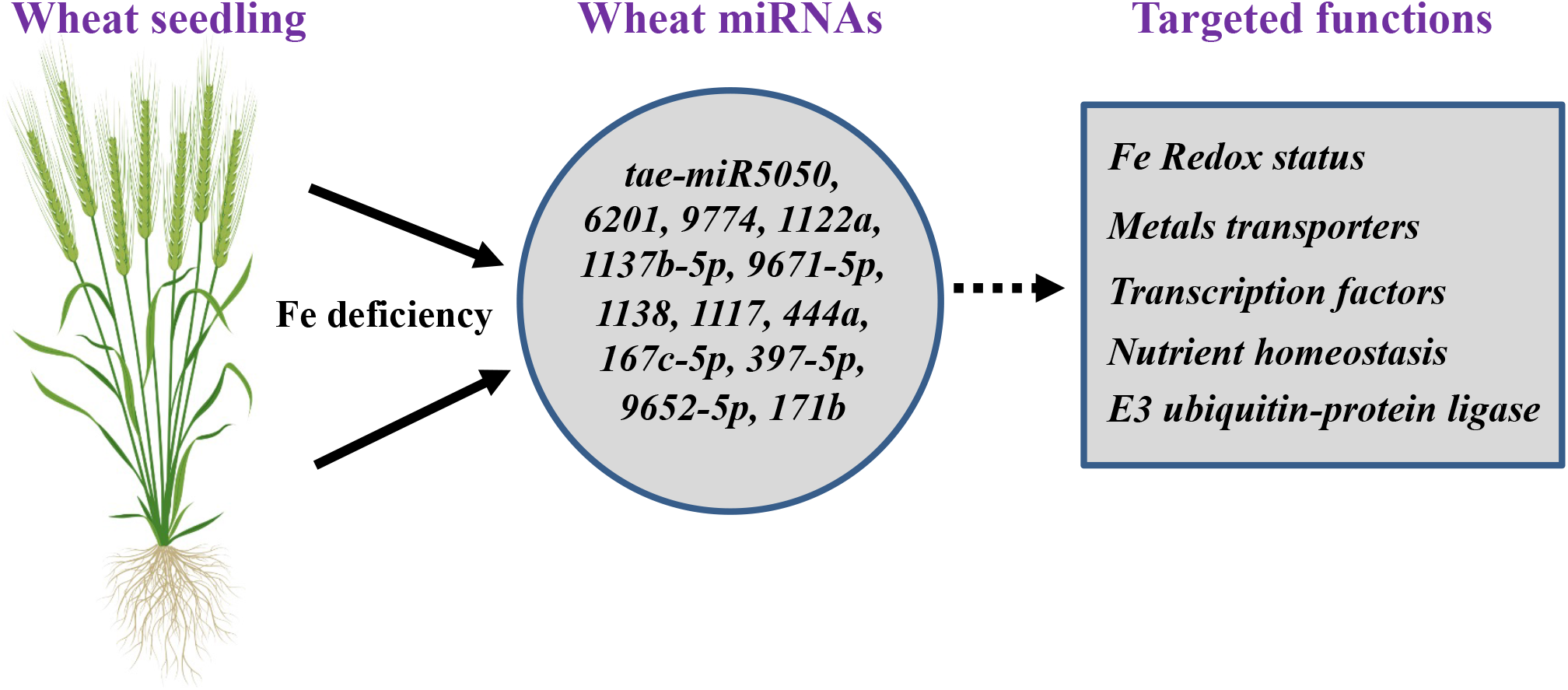
A speculated model for miRNA-mediated Fe homeostasis in hexaploid wheat. The model represents the multiple miRNAs that were DE expressed under Fe deficiency. These miRNA targets distinct genes involved in different biological functions, as mentioned. Some major functions include redox-related metal transporters, transcription factors and E3 ubiquitin ligases.

### Target prediction of wheat miRNAs revealed an adaptive response against Fe deficiency

To identify the regulatory network involved in Fe-homeostasis by these distinct miRNAs, we predicted their targets and analysed them for their involvement in Fe deficiency response. Among the most interesting targets for these Fe deficiency responsive miRNAs revealed a complex network involving multicopper oxidases (MCOs), transcription factors like GRAS and MADS-box, major facilitator superfamily (MFS) transporters, E3 ubiquitin-protein ligases, oxidoreductases, protein kinases etc.. In prediction analysis, we observed that miRNA397-5p could target transcripts encoding for MCOs. *Tae-miRNA171b* was predicted to target transcripts encoding for GRAS TF. On similar lines, sulphate transporters are predicted to be targeted by Fe deficiency-induced miRNA395a and 395b. As we proposed to understand the miRNAs mediated regulatory network against Fe deficiency, we predicted the differential response of shoot and root-expressed miRNAs in terms of their targets. Interestingly, we observed that among the targets for root-induced miRNAs, GRAS family TFs, NAC and MADS TFs, SCR-like genes, serine/threonine phosphatases, and sugar transporters.

While among the targets of root down-regulated miRNAs included potassium transporter, CNX (molecular chaperon), bHLH TFs, MFS transporters, Zn finger TFs, F-box related, sulphate transporters, kinases, cell wall-related genes. Our analysis supported our previous report suggesting over-accumulation of bHLH TFs during Fe deficiency [36]. miRNA-regulated expression of CNX type of molecular chaperones suggested an adaptive response for root against the detrimental effect of ROS accumulation during Fe deficiency. Apart from it, we observed that a significantly downregulated miRNA in the root (*tae-miR 5049-3p*) was found to target *S-adenosyl-L-methionine-dependent methyltransferase* while the S-adenosyl-methionine is the precursor for mugineic acid (MAs) family of PS [41]. This indicates that PS biosynthesis in wheat might be regulated through tae-miR 5049-3p. In the shoot, however, we observed that targets associated with redox enzymes, kinases and phosphatases were over-represented. Among the TFs families, NAC and myb were among the targets of Fe-responsive miRNAs and phytohormone (like auxin and JA) associated genes were also targeted through these miRNAs. Therefore, the target prediction analysis, on the one hand, strengthens the involvement of miRNAs in Fe deficiency response in wheat. At the same time, it also suggests a tissue-specific regulation of their targets.

## Discussion

Earlier miRNA-mediated regulation has been reported under different abiotic and biotic stresses [42–46]. Reports on miRNAs regulating plant adaptation to Fe deficiency and their functional analysis are limited mainly to the model plant *Arabidopsis* [27]. Exploring the miRNA has provided information on the network associations that help crop plants to adapt to different abiotic stresses. However, an attempt has yet to be made to unravel the miRNA-mediated control of Fe homeostasis in hexaploid wheat. Although genes involved in Fe homeostasis were reported earlier, the miRNA-mediated targets were not addressed in wheat [36]. Noteworthy, such molecular responses largely depend on the genotype and specific stress condition. This study attempts to identify wheat miRNA that could be linked to the target pathway functions to identify the critical regulatory miRNA-mRNA interaction involved in Fe deficiency conditions. A total of 105 miRNAs were identified in shoots and roots, respectively. Our work identified 9 novel miRNAs with distinct expression responses during Fe deficiency with a typical stem-loop structure (**Figure S2**).

In this study, sRNA libraries were generated from the roots and shoots of wheat seedlings subjected to Fe deficiency for different time points. This was done to generate the inventory of miRNA that could assist in collating miRNA that may differentially express at any time. Our analysis resulted in the identification of multiple root and shoot specific miRNAs in response to Fe deficiency (Figure 2; Table S3). Among these differentially expressed miRNAs, a subset of randomly selected miRNAs from root and shoot were employed to validate the RNA-Seq data. qRT-PCR of these tissue specific miRNAs drew a strong correlation with respect to RNA-Seq data.. Furthermore, the characterization of differentially expressed miRNAs from root (*tae-miR1138*, -*167c-5p*, -444a, -*9652-5p*, - *9654a-3p* and -*397-5p*) and shoot (*tae-miR6201, -5050, -9774, -1122a, -1137b-5p* and *-9671-5p*) revealed the spatio-temporal expression of these Fe-responsive miRNAs.

Fe deficiency is a major nutritional disorder that limits crop productivity. In plants, multiple miRNA gene families are known to be involved in the Fe deficiency responses [27,47]. In addition, correlation studies were done where the expression of specific miRNA was observed in high and low Fe genotypes of wheat and rice [30,48]. These studies support that miRNA-mediated control could occur concerning the Fe flux in a tissue-specific manner. In our analysis size of the majority of the filtered reads was 21-24 nt (**Figure 1E**). Specifically, the 24 nt size class represented the highest among the total sizes. This range is consistent with the previous reports from other plant species [49–51].

Wheat DE miRNA targets candidate genes from the family of TFs, such as MADS, GRAS, WRKY, and F-box-containing proteins. At the transcript level, miRNA targets include ferroxidases, E3 ubiquitin ligases and enzymatic reaction associated genes. Previous results have shown that members of the above TFs and other metabolic genes showed up-and/or down-regulation during Fe deficiency [27,36]. This suggests that miRNA responding to the Fe deficiency targets Fe-responsive genes to modulate nutrient homeostasis. Interestingly, one of the primary targets of this study identified encoding multiple MCOs belonging to the laccases gene family. Wheat recruit strategy-II mode of Fe^3+^ in its chelated form. The generation of the Fe^3+^ in the periplasmic space is primarily controlled by the ferroxidase activity of the laccases subfamily [52,53]. In plants, multiple MCOs were shown to be differentially expressed under Fe deficiency [54]. Consistant to these studies, in our analysis *tae-miR397-5p* shows potential targets for multiple wheat MCOs **(Table S6)**. This suggested a mechanistic insight that regulated the turnover of wheat ferroxidase activity in wheat through *miR397* at the post-transcriptional level. Earlier studies demonstrated the *Arabidopsis miR397* role in imparting improved plant tolerance to cold stress [55]. In barley, the expression of *miR397* decreased under the Boron toxicity [56] and citrus plants [31]. The expanded role of *miR397* suggests its involvement in enhancing the rice grain size by promoting panicle branching [57]. Another candidate, *tae-miR444*, was highly upregulated in Fe deficiency. The monocot-specific rice *miR444* was shown to be an important factor in relaying the antiviral signalling from virus infection to plant RNA-dependent RNA polymerase1 [58]. At the mechanistic level, it was shown that *miR444* could repress the MADS-box encoding transcript. Our target prediction suggests that *Tae-miR444* could target multiple wheat MADS-box and F-box containing encoded transcripts. MADS-box encoded transcripts are differentially regulated during Fe deficiency conditions. *Tae-miR1122* also shows root-specific induction in Fe deficiency. Based on the prediction searches, *Tae-miR1122* was identified to be differentially expressed in the EST libraries represented by cold stress and Aluminium exposed wheat root tips and seedlings [59]. Earlier reports have suggested that expression of miRNAs under different nutrient deficiencies influences the adaptation to different conditions. For example, *Arabidopsis* miRNAs show overlapping responses during multiple nutrient deficiency conditions. This suggests the multifunctional role of miRNA that may be commonly upregulated or suppressed by certain nutrients [60].

Research on sulphur (S) and Fe interaction suggests that Fe uptake depends on the assimilation of the S in the roots [61-63]. S in the soil is absorbed as SO_4_^2−^ in root tissue and is translocated to the aerial parts in its reduced forms to be utilized in the subcellular organs such as chloroplasts and mitochondria for cysteine and methionine biosynthesis. Strategy II plants primarily rely on releasing PS in the rhizosphere to mobilise soil Fe. The slow rate of PS biosynthesis and decreased nicotianamine (NA) level in the cells were linked to the S deficiency. Therefore, sulphate metabolism and plant distribution have been linked with Fe uptake and translocation. Plant sulphate transporters (SULTR) were differentially regulated under the changing regimes of Fe [36,64]. A S-containing compound such as glutathione (GSH) shows a link with the tolerance to Fe [65]. Therefore, the differential regulation of genes involved in sulphate uptake and its metabolism should be tightly regulated. miR395 is an integral part of sulphate assimilation that regulates the expression of SULTR to maintain sulphate uptake and utilization in plant tissue. Fe deficiency-induced wheat miRNA395 supports the notion of the cross-talk regulation between the Fe and S homeostasis in monocots. Our study observed high target scores for multiple SULTR targeted by wheat miR395 (Supplementary Table S4). We conclude that Fe deficiency-induced recruitment of post-transcriptional regulation could affect the secondary process regulating nutrient uptake.

Comparison of miRNA repertoires between wheat and its diploid progenitors provides useful information about the changes in miRNA gene content over time and the role of miRNAs in wheat’s adaptation to its environment [66]. Our study observed a high proportion of sRNA contribution from A and B genomes compared to D genomes. This was consistent for both +Fe and -Fe datasets. Genome expression bias under stress conditions at the transcriptome level (mRNA) has been reported earlier in hexaploid wheat with high contribution from the A genome [67, 68, 36, 38]. Although analysis of such biasness for miRNA is tedious, we tend to calculate the contribution of the miRNA putatively arising from different genomes. Our analysis for the –Fe regulated miRNAs points to a high expression contribution in the ancestral genotypes of wheat with the highest for the *T. aestivum* (AABBDD) and *T. durum* c.v. Langdon TTR16 (AABB) compared to *Aegilops tauschii* (DD genome). Our observation agrees with previous reports, suggesting the least involvement of DD genome in miRNA diversity [66]. It could be possible that incorporating the DD genome into the AABB genome increased the expression of miRNAs, suggesting a synergistic effect or some trans-genomic regulation. The reasoning for these observations could be answered by extended miRNA genome bias expression that remains to be investigated.

Altogether, miRNA profiling suggests their involvement in regulating Fe deficiency responses. Our work provides evidence that miRNA perturbed due to Fe deficiency targets a subset of previously reported Fe-responsive genes (**Figure 7**), This reflects that miRNA-mediated control of Fe-responsive genes contributes to such regulatory mechanism in hexaploid wheat. Specifically, the miRNA can target the genes primarily involved in changing the Fe redox status and its uptake. Similarly, the regulation of multiple TFs was also targeted by miRNA (**Figure 7**). Overall, the generated datasets will serve as an important resource to further investigate the transcriptional rearrangements that influence different tiers of molecular response during Fe deficiency. A further study focusing on the candidate miRNAs function could add a new paradigm into Fe deficiency and other stress to improve plant growth and yield.

## Material and methods

### Plant material, Fe deficiency treatment and sampling

Hexaploid wheat cv. “C-306” was used for this experiment. Wheat grains were stratified in the dark at 4°C overnight and germinated for 4 days on Petri plates lined with Whatman filter paper. The endosperms were removed from the developing seedlings to cut out the nutrient supply from the seed. Subsequently, the seedlings were grown for 5d in Hoagland’s nutrient solution and then subjected to Fe deficiency treatment. For Fe starvation (–Fe), 2 µM Fe (III) EDTA was used as the Fe source. For control plants (+Fe), Hoagland’s nutrient solution was used, keeping other nutrients unchanged with 80 µM Fe(III) EDTA. Plants were grown in the growth chamber set at 21±1°C, 50–65% relative humidity, and a photon rate of 300 μmol quanta m^−2^ s^−1^ with a 16 h day/8 h night cycle. For sampling, roots, and shoots were collected from three biological replicates at different time points, i.e., 6, 9, 12 and 15 days after deficiency, and crushed in liquid nitrogen. Total RNA extraction was performed from the root and shoot at the indicated time points (**Figure S1**). RNA samples from these time points were pooled in equal proportions for the control and Fe deficiency samples of root and shoot tissue, respectively.

### Preparation of Small RNA library and RNA sequencing

The total RNA of control and treated roots and shoots was extracted using Trizol Reagent (Invitrogen, ThermoFisher, USA) according to the manufacturer’s instructions. The RNA quantity and purity were assessed using NanoDrop™ One (ThermoFisher Scientific) and denatured agarose gel electrophoresis. RNA integrity number (RIN) was checked using Bioanalyzer 2100 (Agilent Technologies, USA), and samples proceeded for small RNA library preparation. The library was constructed by TruSeq Small RNA Sample Prep Kit (Illumina, USA). Small RNAs were ligated with 3’ and 5’ adapters, followed by reverse transcription, PCR enrichment, purification, and size selection. The sRNA libraries were sequenced on the NovaSeq6000 Illumina Sequencing platform. Transcriptome sequencing data were deposited at NCBI (Submission: SUB12485490 with BioProject: PRJNA916207).

### Bioinformatics analysis and miRNA identification

The data obtained from high throughput sequencing was processed into raw sequencing reads by CASAVA base recognition. Low-quality raw reads containing adaptors, 5’ primer contamination, and polynucleotide tails, and reads with >50% bases having a Qphred less than or equal to 5, and the ones in which >10% base information were indeterminable were discarded. The clean reads ranging from 18 to 30 nucleotides were mapped to the reference genome sequence of *T. aestivum* using Bowtie to analyze their expression level and distribution on the genome. Rfam database and Repeat Masker were used to remove non-coding RNAs – rRNA, tRNA, snRNA, snoRNA, and repeat sequences. The unmatched reads were classified as putative miRNAs and subsequently aligned against miRbase (http://www.mirbase.org/) to obtain detailed information on mapped reads, including the secondary structure of mapped miRNAs, the sequence of miRNAs in each sample, their length, and occurrences. Matched sequences were identified as conserved miRNAs, and the characteristic hairpin structure of other remaining miRNAs (marked as novel miRNAs) was predicted by miRDeep2, miREvo software and RNA fold server (http://rna.tbi.univie.ac.at/cgi-bin/RNAWebSuite/RNAfold.cgi) [69,70].

### Identification of differential expressed miRNAs

To investigate the differentially expressed miRNAs between (+)Fe or (–Fe) libraries, the expression of known and unique miRNAs in each sample was statistically analysed by transcripts per million (TPM) [71]. Read counts were normalized to TPM as follows: The normalized expression = (read count*1,000,000)/Mapped reads. The differential expression level of miRNAs was calculated using DEGseq [72] and miRNAs with log2 fold change >1 and p-value < 0.01 were considered as differentially expressed.

### miRNA target prediction and annotation

Wheat target genes for known and novel microRNAs were predicted using the psRobot and psRNATarget tool with default parameters [73,74]. To determine the functional categorization of miRNA target genes, Gene Ontology analysis was carried out using PANTHER classification system (http://pantherdb.org/) [75]. p-values were corrected using the Bonferroni method, and GO terms with adjusted p-value <= 0.05 were considered significantly enriched. GO terms enriched by more than 3 folds were plotted using the ggplot2 package from R studio [76]. To gain insight into the metabolic pathways of target genes, KEGG pathway enrichment was carried out by KOBAS software [77]. Owing to lack of wheat dataset in KOBAS, rice dataset was used to identify significantly enriched KEGG pathways [36,78]. We annotated the wheat sequences with RefSeq rice dataset using BLASTN and cut-off score 1e-10, and used adjusted p-value of 0.05 to obtain the significant pathways.

### Quantification of miRNAs by real time-PCR

To get an insight into the spatio-temporal expression patterns of differentially expressed miRNAs and to validate the transcriptome data, stem-loop qRT PCR analysis was conducted. Primers were designed following the method of [79] and are listed in **Table S5**. Briefly, 1μg of DNase-treated total RNA was reverse-transcribed using TaqMan microRNA reverse transcription kit (Applied Biosystems™) according to the manufacturer’s instructions. The real-time PCR program was set as follows: 95°C for 3 min, 45 cycles (95°C for 10 sec, 55°C for 20 sec,72°C for 20 sec). All reactions were performed in triplicate for each time point. The relative expression levels of the miRNAs were calculated by the 2^-^ΔΔCT method [80]. For normalization, the wheat gene U6 (GenBank: X63066.1) was used as an internal control [81].

All qRT-PCR was performed using SYBR Green I (Takara SYBR Premix Ex Taq) on the Bio-Rad CFX96 Real-time PCR detection system.

The spatiotemporally pooled total RNA samples of Fe sufficient and deficient root and shoot tissues (individually) used for transcriptomic analysis were employed for qRT-PCR-based validation of the RNA-Seq data. Eight miRNAs were randomly selected to test the correlation between the two data sets.

### Expression analysis of differentially expressed miRNAs in wheat database

For expression analysis of differentially expressed miRNAs in parent cultivars of wheat, we utilized our in-house Plants miRNA expression atlas database (PmiRExAt, http://pmirexat.nabi.res.in/). The expression from the parent lines was collated for the whole plant, including *T. aestivum* (AABBDD), *T. durum* c.v. Langdon TTR16 (AABB) and *Aegilops tauschii*, TQ113 (DD). The table was collated using the TPM values and comparative analysis was done using pie charts. The contribution of different subgenomes was calculated based on the number of miRNAs detected in any specific genome (penetrance) and the level of miRNA expression in any particular subgenome (expressivity).

## Availability of data and materials

The small RNA-Seq raw read data has been submitted at NCBI (Submission: SUB12485490 with BioProject: PRJNA916207).

## Acknowledgments

The authors thank the Executive Director of NABI for the facilities and support. This work was funded by the Department of Science and Technology-Science & Engineering Research Board (SERB) Grant number CRG/2020/000940 and the NABI-CORE grant to AKP. DBT-eLibrary Consortium (DeLCON) is acknowledged for providing timely support and access to e-resources for this work. The wheat genome resources developed by International Wheat Genome Sequencing Consortium are highly appreciated.

## Author contribution

AKP and SS conceived and designed the research. SS and APS carried out wet lab experiments; SS, APS, RJ, PS, DS, SM and VS carried out wet lab experiments and performed data analyses. AKP, SS, APS and DS wrote and finalized the manuscript. All authors have read and approved the final manuscript.

## Ethical approval

Not applicable.

## Consent for publication

Not applicable.

## Supplementary Figures

**Figure S1.The schematic** diagram for workflow pipeline to identify the small RNAs and development of miRNA inventory.

**Figure S2.Small RNA reads distribution per chromosome of the wheat genome.** Images showing a graphical representation of sRNA reads for **A)** Control shoot, **B)** Fe deficient shoot, **C)** Control root, and **D)** Fe deficient root. The chromosome is shown as the outer circle. Grey background in the middle area shows the distribution of 10,000 reads on the chromosome. Red in the gray background represents the number of sRNAs on the sense strand of the chromosome, while blue represents the number of sRNAs on the antisense strand. All reads are shown in the center area of the circle, where red represents the number of sRNAs on the sense strand of the chromosome, and the green represents the number of sRNAs on the antisense strand.

**Figure S3.Secondary structure plots of novel identified miRNAs.** The hairpin structure of each novel miRNA was predicated with RNA fold server (http://rna.tbi.univie.ac.at/cgi-bin/RNAWebSuite/RNAfold.cgi) using the pre-miRNA sequence based on minimum free energy.

**Figure S4.Genome biased expression of miRNAs in wheat.A)** Heat map was generated using the PmiRExAt http://pmirexat.nabi.res.inserver. Values in the heat map represent the TPM (transcripts per million) values. **B)** Average number of miRNAs being expressed in different wheat genomes. **C)** Average relative expression of miRNAs in different wheat genomes.

## Supplementary Tables

**Table S1.** Small RNAs obtained in Control (Fe-EDTA 80µM) and –Fe treated (Fe-EDTA 2µM) Wheat C-306 small RNA libraries of root and shoot

**Table S2.**Positional mapping of wheat small RNAs to exon and intron

**Table S3.**List of differentially expressed miRNAs in response to Fe deficiency, along with their precursor and mature sequences. Values against each miRNA indicate the log_2_ fold change observed in the expression.

**Table S4.**Hairpin family classification of all the miRNAs identified in this study and across different plant species. “+” means that the miRNA family exists in this species, and a “-” means the miRNA family does not exist in this species.

**Table S5.**Primer sequences of the DE miRNAs for expression analysis by stem-loop qRT-PCR.

**Table S6.**List of target genes for DE miRNAs in hexaploid wheat roots and shoots identified by pSRNATarget.

